# Paternal over- and under-nutrition program fetal and placental development in a sex-specific manner in mice

**DOI:** 10.1101/2025.11.14.688439

**Authors:** Hannah L. Morgan, Nader Eid, Nadine Holmes, Matthew Carlile, Sonal Henson, Fei Sang, Victoria Wright, Marcos Castellanos-Uribe, Iqbal Khan, Nazia Nazar, Sean T. May, Rod T. Mitchell, Federica Lopes, Robert S. Robinson, A. Augusto Coppi, Vipul Batra, Adam J. Watkins

## Abstract

The association between sub-optimal paternal diet and offspring well-being is becoming established. However, the underlying mechanisms are yet to be fully defined. The aim of this study was to establish the impact of over- and under-nutrition, with or without macronutrient supplementation, on male reproductive fitness and post-fertilisation development. Male C57/BL6J mice were fed either control diet (CD), isocaloric low protein diet (LPD), high fat/sugar ‘Western’ diet (WD) or LPD or WD supplemented with methyl-donors and carriers (MD-LPD or MD-WD respectively) for 8 weeks before mating with virgin C57/BL6J females. Placental tissue was collected at embryonic day (E)8.5, to assess early placental (ectoplacental cone) morphology and metabolism and E17.5 for sex-specific transcriptomic profiling. Post-mating, stud male tissues were harvested for assessment of testicular morphology and gene expression, gut microbiota composition and metabolic status. WD and MD-WD males displayed increased adiposity, hepatic cholesterol and free fatty acids and gut microbiota dysbiosis when compared to CD fed males. In the testes, WD and MD-WD perturbed the expression of genes associated with metabolism and transcription regulation. Additionally, we observed differential expression of multiple genes within the Wnt signalling pathway, central in the regulation of cellular proliferation, migration, survival, and cell fate determination during development. Despite no impact on fundamental male fertility, significant changes in ectoplacental cone metabolism, fetal growth, and placental gene expression were observed in response to specific dietary regimens. Interestingly, while CD male and female placentas displayed 301 genome-wide, sexually-dimorphic genes, LPD, MD-LPD, WD and MD-WD male and female placentas possessed only 13, 0, 14 and 15 sexually-dimorphic genes respectively. Our data show that while sub-optimal paternal diet has minimal impact on male fertility, fetal and placental development are perturbed in a sex-specific manner.

## Introduction

Under- and over-nutrition are directly linked to poor physiological, metabolic and reproductive health. The prevalence of poor dietary habits is increasing worldwide, with both under- and over-nutrition rising dramatically. Current estimates suggest that the combined prevalence of under- and overweight has increased in over 160 countries in both adults and school-age children (Collaboration, 2024). Therefore, there has been significant interest in the impact of sub-optimal nutrition and dietary supplementation on physiological, metabolic, and reproductive health (Jawaid et al., 2021).

The consumption of diets high in fat and/or sugar are central to the development of various metabolic disorders including obesity, hypertension, non-alcoholic fatty liver disease (NAFLD; also referred to as Metabolic Dysfunction-Associated Steatotic Liver Disease, MASLD) (Ludwig et al., 2001, Nielsen et al., 2010, von Frankenberg et al., 2017) and metabolic syndrome (O’Neill and O’Driscoll, 2015, Saklayen, 2018). Poor quality diet has also been linked to impairments in fertility and reproductive fitness. Obesity has been linked to a reduction in testosterone levels, sperm concentration, motility and viability, and damage to the seminiferous tubules of the testes (Crean et al., 2023, Fan et al., 2015, Leisegang et al., 2014, Lotti et al., 2013). Undernutrition has also been linked to multiple metabolic and reproductive impairments. In rodents, a low protein diet (LPD) significantly reduces body weight, weight of the testes, epididymis and seminal vesicles, and serum testosterone and follicle-stimulating hormone levels (Ajuogu et al., 2020).

Poor paternal diet also impacts negatively on fetal development and adult offspring health (Batra et al., 2022, Capobianco and Pirrone, 2023, Fleming et al., 2018). Impairments in rodent fetal and placental development have been reported in response to diets low in protein (Morgan et al., 2021, Watkins et al., 2017), folate (Lambrot et al., 2013) and high in fat. (Jazwiec et al., 2022). Separately, children born to fathers with early onset type-2 diabetes are leaner and have a higher insulin sensitivity than children from healthy men (Penesova et al., 2010). Additionally, children from men with type-2 diabetes at the time of conception have a lower weight at birth, but with increased BMI in later life and a greater risk of developing type-2 diabetes and hyperlipidaemia themselves (Moss and Harris, 2015, Penesova et al., 2010, Silva et al., 2017). In rodent models, paternal high fat and sugar diets perturb offspring insulin and glucose homeostasis (Cesar et al., 2022), behaviour and gut microbiota composition (Bodden et al., 2022). Interestingly, poor paternal diet impacts on offspring health in a sex-specific manner. Sex-specific changes in offspring tissue lipid abundance have been reported following a paternal low protein diet (Furse et al., 2023, Morgan et al., 2022, Morgan et al., 2020b, Watkins et al., 2018). Similarly, paternal high fat diet in rats programs pancreatic β-cell dysfunction and impaired glucose metabolism in female offspring (Ng et al., 2010) and adult-onset obesity in males (Sanchez-Garrido et al., 2018).

Separate to macro-nutrient modification, the significance of dietary micro-nutrients such as vitamins, minerals and antioxidants has also been assessed. Studies have focused on supplementation as a potential means to reverse or alleviate the effects of sub-optimal diets. In rats, the detrimental effects of a high fat/sucrose diet on liver steatosis were ameliorated through the supplementation with a range of methyl donors and carriers such as folate, choline, betaine, and vitamin B12 (Cordero et al., 2013a, Cordero et al., 2013b) In mice, supplementing calorically restricted males with vitamins and antioxidants prevented sperm oxidative damage and normalized offspring growth and adiposity (McPherson et al., 2016). Similarly, supplementation of a paternal LPD with methyl donors and carriers has been shown to normalise patterns of fetal over-growth in mice (Morgan et al., 2020a).

While the association between paternal diet and offspring well-being is becoming established, the underlying mechanisms are yet to be fully defined. The aim of this study was to establish and compare the impact of over- and under-nutrition on male reproductive fitness and post-fertilisation development. As poor micronutrient status has also been associated with perturbed fetal development (Lambrot et al., 2013), we additionally explored the role that supplementation with specific methyl donors and carriers may play in ameliorating any detrimental effects of poor paternal diet.

## Results

### Male physiology and metabolic status

To investigate the impact of over- and under-nutrition, with or without methyl-donor and carrier supplementation, on male reproductive fitness, we fed male C57BL/6J mice one of five diets comprising a control diet (CD), low protein diet (LPD), LPD supplemented with methyl donors (MD-LPD), Western diet (WD) or WD supplemented with methyl donors (MD-WD) for up to 24 weeks. Over the course of the feeding period, all experimental groups became lighter than CD males (Figure 1A). However, no significant differences were observed when analysed at individual time points or as average growth profile (area under the growth curve). At the time of cull, there was no difference in mean testicular (Figure 1B) or seminal vesicle (Figure 1C) weights. MD-WD had an increased total fat mass (combined weight of gonadal, peri-renal, inguinal and interscapular fat pads) when compared to CD fed males (Figure 1D, P = 0.01). Analysis of white (WAT, combined weight of gonadal, perirenal and inguinal fat) to brown adipose tissue (BAT; interscapular) ratio revealed an increase in WD and MW-WD males when compared to CD fed males (Figure 1E, P <0.05). Next, we assessed serum profiles of central indicators of inflammatory status (Tnf) and testicular regulation (Inhibin β-A chain). There was no difference in the concentrations of Tnf (Figure 1F) between males. However, WD fed males displayed an elevated concentration of serum Inhibin β-A chain, a paracrine regulator of germ cell proliferation, spermatogenesis and Leydig cell steroidogenesis, when compared to LPD and MD-LPD fed males (Figure 1G, P <0.01).

**Figure 1.**
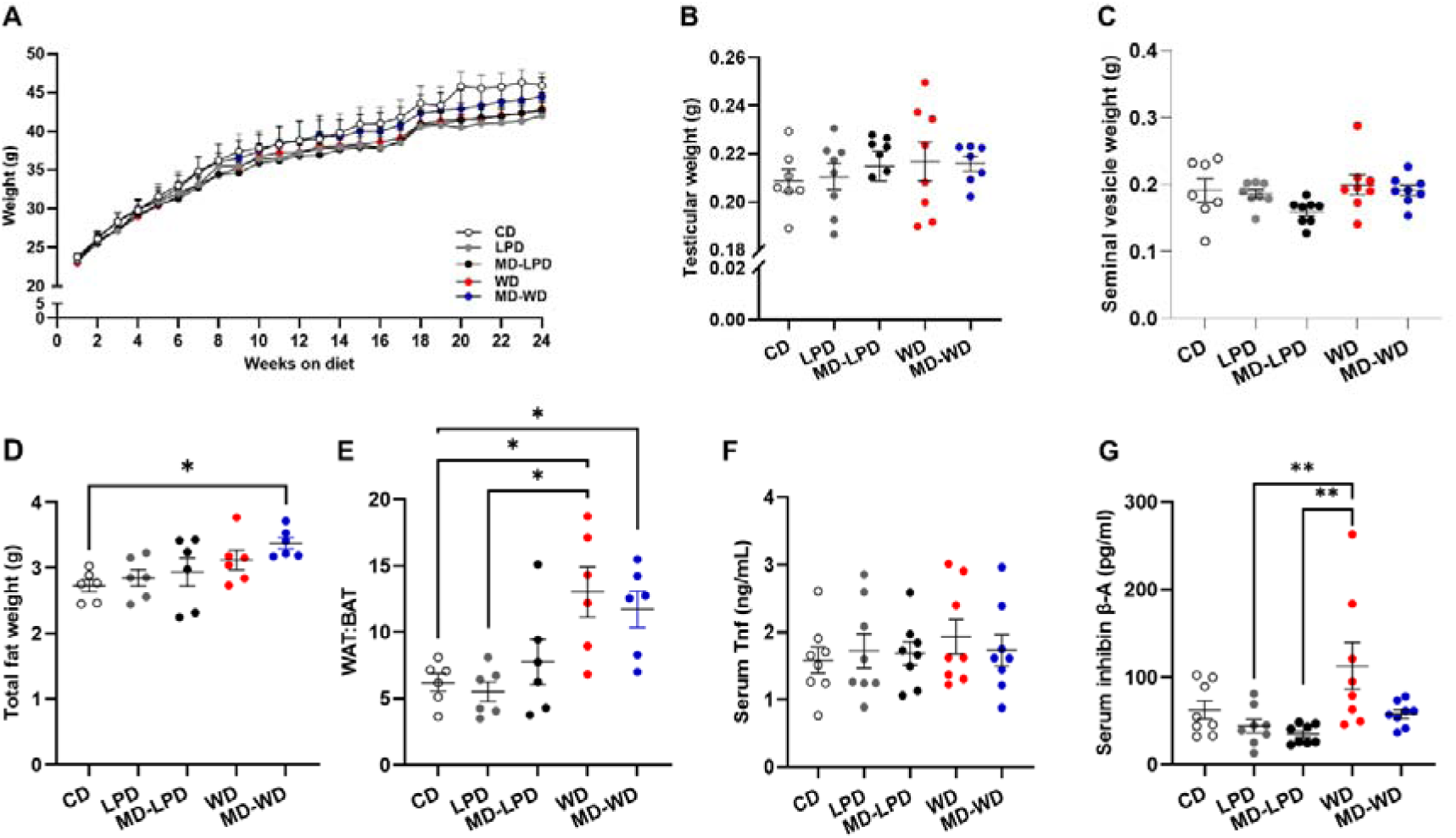
Impact of diet on male growth. (A) Mean weekly body weight of males fed either a control diet (CD), low protein diet (LPD), methyl donor-supplemented LPD (MD-LPD), Western diet (WD) or methyl donor-supplemented Western Diet (MD-WD). Mean (B) testicular weight, (C) seminal vesicle weight (D) total fat weight (combined weight of individual fat pads) and (E) ratio of white adipose tissue (WAT) to brown adipose tissue (BAT). Mean serum (F) Tnf and (G) inhibin β-A chain levels. N = 6-8 males in each group. Data were analysed using a one-way ANOVA with Holm-Sidak post hoc tests for multiple comparison. * P <0.05, ** P <0.01.

As we observed significant changes in adiposity in response to our diets, we assessed stud male metabolic status. There were no differences in the concentrations of serum glucose (Figure 2A) or insulin (Figure 2B) between groups. In contrast, WD males displayed an elevated concentration of hepatic cholesterol when compared to CD males (Fig, 2C) while MD-WD males displayed elevated levels when compared to CD, LPD and MD-LPD males (Fig, 2C, P <0.05). WD males also displayed elevated levels of free fatty acids (FFAs) when compared to CD, LPD and MD-LPD males (Figure 2D, P <0.05), while the levels of FFA in MD-WD males was not significantly different to that of either CD or WD males. No differences in the concentration of hepatic triglyceride were observed between groups (Figure 2E).

**Figure 2.**
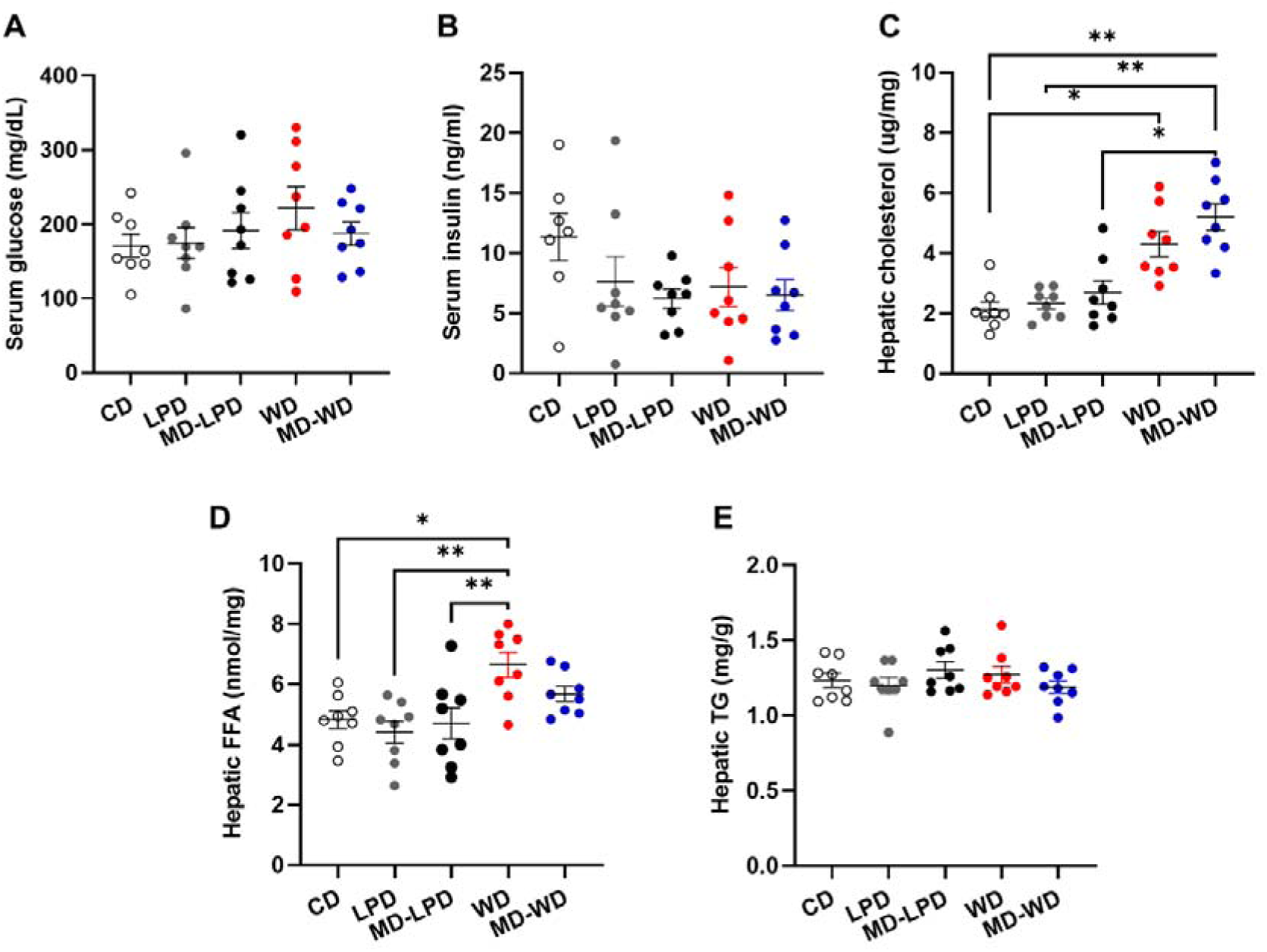
Impact of diet on male non-fasting metabolic status. Serum (A) glucose and (B) insulin in males fed either a control diet (CD), low protein diet (LPD), methyl donor-supplemented LPD diet (MD-LPD), Western diet (WD) or methyl donor-supplemented WD (MD-WD). Hepatic (C) cholesterol, (D) free fatty acids (FFAs) and (E) triglyceride (TG) concentrations. N = 8 males in each group. Data were analysed using a one-way ANOVA with Holm-Sidak post hoc tests for multiple comparison. * P <0.05, ** P <0.01.

Analysis of gut bacterial profiles revealed no difference in overall bacterial diversity (Figure 3A) or species evenness (Figure 3B) between groups. When analysed at the phylum level (Figure 3C), MD-WD males displayed a significant increase in the abundance of Deferribacteres (Figure 3D; P = 0.042), while WD and MD-WD males displayed an increased abundance of Proteobacteria (Figure 3E; P <0.05) when compared to CD males. Furthermore, the abundance of TM7 (Saccharibacteria) was decreased in MD-WD males when compared to LPD and MD-LPD fed males (Figure 3F; P <0.05). Analysis of bacteria at the family level identified significant reductions in the abundance of Turicibacteraceae and Lachnospiraceae in both WD and MD-WD males when compared to CD fed males (Figure 3G, P <0.05). Additionally, the abundance of S24-7 (Muribaculaceae), Clostridiaceae, Dehalobacteriaceae, Ruminococcaceae and Desulfovibrionaceae were all significantly altered (both increased and decreased) in MD-WD males when compared to CD males (Figure 3G; P <0.05).

**Figure 3.**
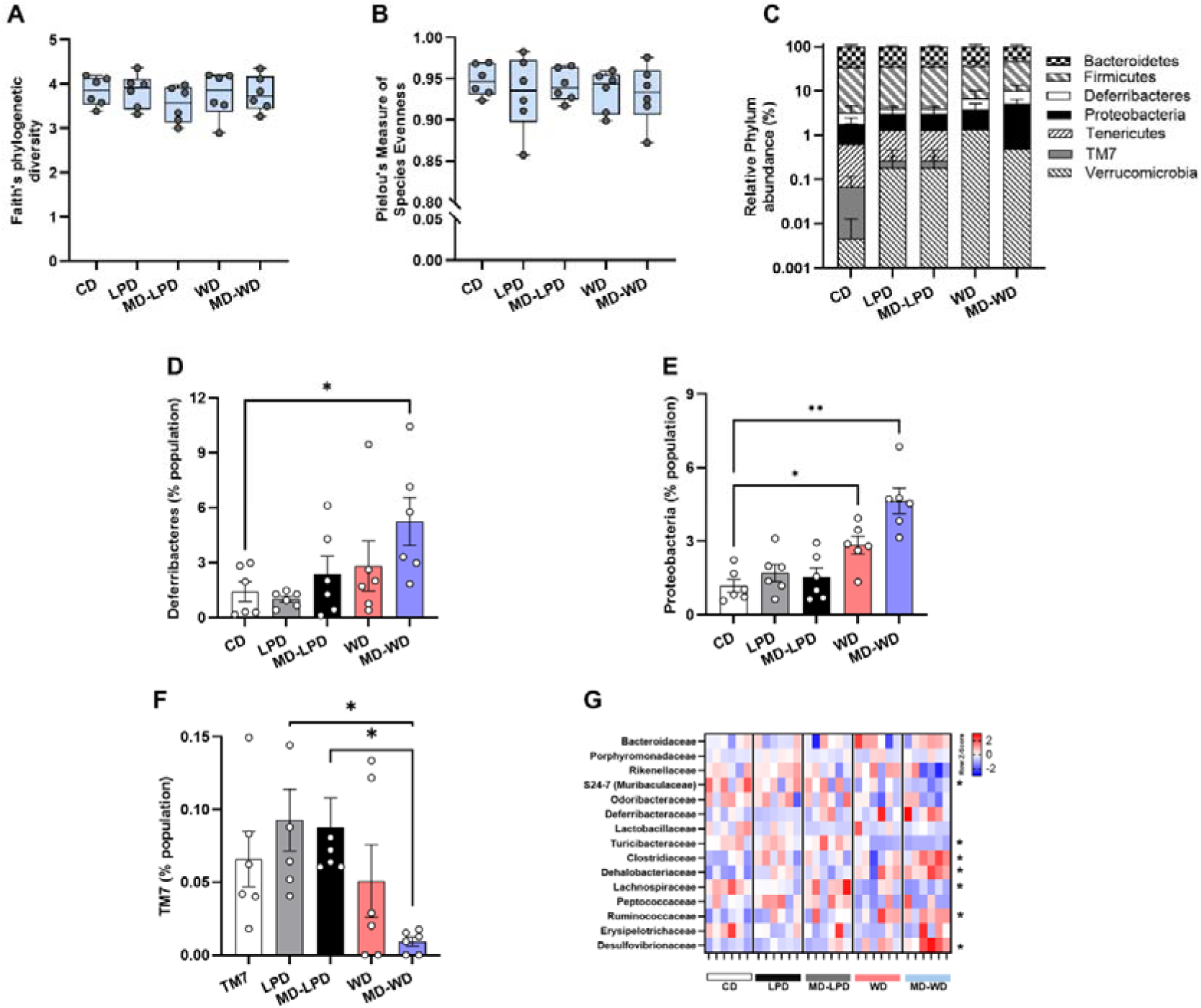
Impact of diet on male gut microbiota. (A) Faith’s phylogenetic diversity and (B) Pielou’s measure of species evenness. (C) Overall bacterial abundance at the phylum level and relative abundance of (D) Deferribacteres, (E) Protobacteria and (F) TM7 phylum. (G) Relative (Z-score) abundance of bacteria at the family level in males fed either a control diet (CD), low protein diet (LPD), methyl donor-supplemented LPD diet (MD-LPD), Western diet (WD) or methyl donor-supplemented WD (MD-WD). N = 8 males in each group. Data were analysed using either a one-way ANOVA (panels D, E, F and G) or Kruskal–Wallis test (panels A, B and C) with Holm-Sidak or Dunn’s post hoc tests for multiple comparison respectively. * P <0.05, ** P <0.01.

### Sub-optimal diet alters seminiferous tubule cytoarchitecture and cellular distribution

As diet and nutritional status have been directly linked to reproductive health, we next assessed testicular and epididymal morphology. Gross histological analysis (Figure 4A) of testicular tissue revealed an increased number of tubules displaying abnormalities such as vacuoles, loss of the epithelium or complete loss of the tubular architecture (Figure 4B) in WD and MD-WD males when compared to CD, LPD and MD-LPD males (Figure 4C, P<0.01). To analyse seminiferous tubule composition further, we stained testicular sections from the same males to determine Sertoli (Sox-9), spermatocytes and spermatids (Ddx4) and spermatogonial stem (Plzf) cells (Figure 4D) numbers. There were no differences in the mean number of nuclei per tubule (as determined via DAPI staining; Figure 4E), the number of Sertoli cells per tubule (Figure 4F) or the number of spermatocytes and spermatids per tubule (Figure 4G). However, a significant reduction in the number of Plzf^+^ spermatogonial stem cells was detected in tubules from LPD and WD males when compared to CD males (Figure 4H, P <0.05). In contrast, no difference in the number of Plzf^+^ cells were observed in MD-LPD and MD-WD testes when compared to CD males (Figure 4H).

**Figure 4.**
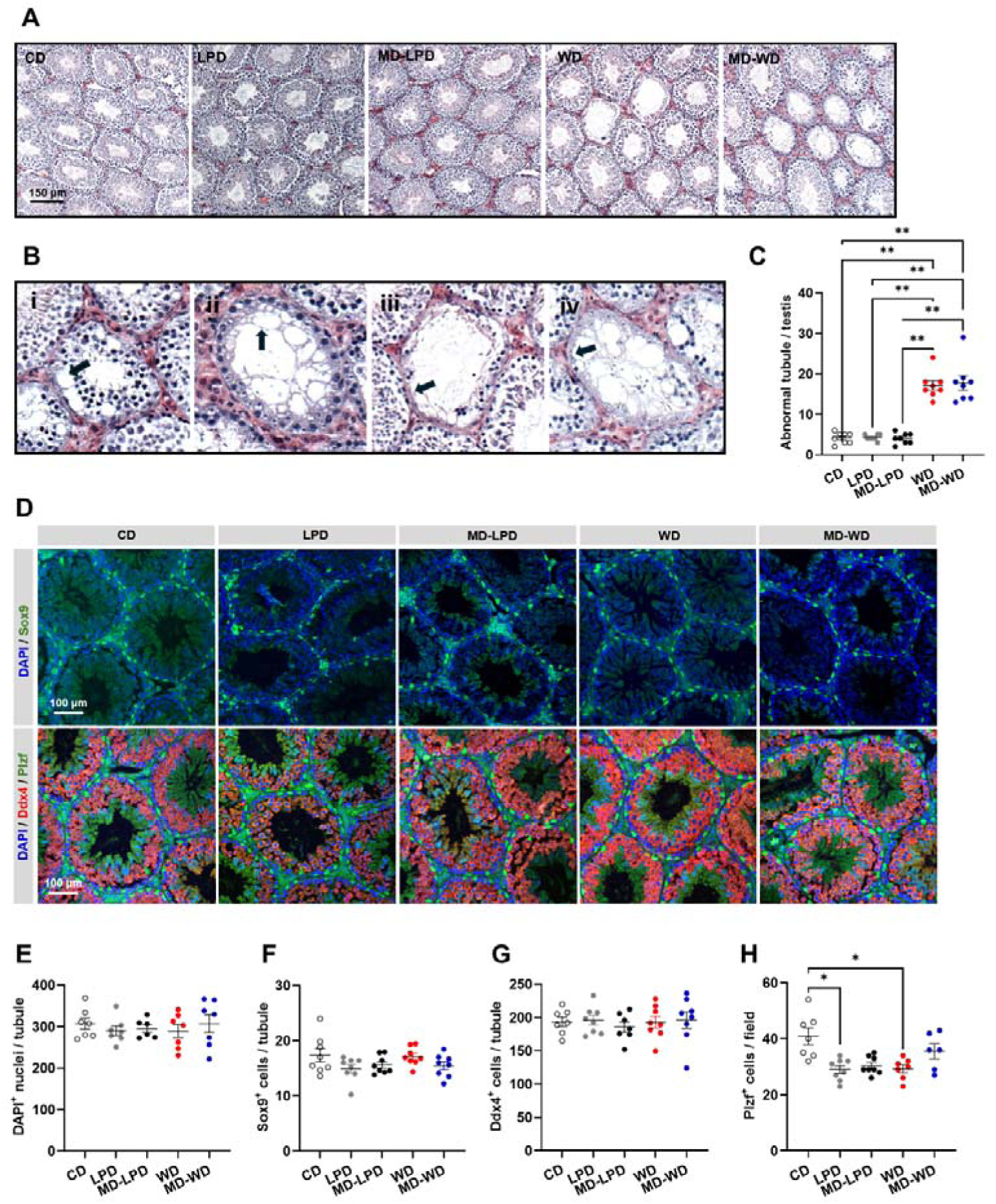
Impact of diet on male testicular morphology. (A) Representative images showing tubule morphology in males fed either a control diet (CD), low protein diet (LPD), methyl donor-supplemented LPD (MD-LPD), Western diet (WD) or methyl donor-supplemented WD (MD-WD). (B) Examples of tubule anomalies identified in WD and MD-WD males including separation of the epithelium from the tubule basement membrane (as indicated by an arrow in i), appearance of vacuoles (as indicated by an arrow in ii), and loss of the germinal epithelium (as indicated by arrows in iii and iv). (C) Frequency of abnormal tubules per testis in males fed either CD, LPD, MD-LPD, WD or MD-WD. (D) Representative staining patterns for DAPI (nuclear counterstain), Sox9 (marker of Sertoli cells), Ddx4 (marker of spermatocytes and spermatids) and Plzf (marker of spermatogonial stem cells) in testes from males fed CD, LPD, MD-LPD, WD and MD-WD. (E) Number of DAPI, (F) Sox9, (G) Ddx4 and (H) Plzf^+^ cells. N = 7-8 males in each group. Data were analysed using either a one-way ANOVA (panels E, F, G and H) or Kruskal–Wallis test (panel C) with Holm-Sidak or Dunn’s post hoc tests for multiple comparison respectively. * P <0.05, ** P< 0.01.

### Testicular gene expression is perturbed in a diet specific manner

To define the impact of diet on testicular gene expression profiles, we performed a micro-array transcriptomic analysis using testicular tissue from the same males. We identified 0, 0, 402 (267 down-regulated, 135 upregulated) and 285 (187 down-regulated, 98 up-regulated) differentially expressed (fold change <-1 or >1; Padj <0.05) genes in LPD, MD-LPD, WD and MD-WD testes when compared to CD testes (Figure 5A-C, F). In WD testes, we identified 267 downregulated and 135 upregulated genes (Figure 5C) associated with an up regulation of fatty acid metabolism (GO:0006631) and a down regulation of chromatin binding (GO:0003682), RNA binding (GO:0003723) and actin filament organization (GO:0007015) pathways (Figure 5D). Network analysis of the 402 differentially expressed genes within the WD testes identified several modular relationships between genes (Figure 5E). Specifically, we observed significant (Padj <0.05) gene nodes associated with the expression of *Ywhae* (FC -1.14), *Rara* (FC -1.35), *Dnmt1 (*FC -1.36), *H3f3a* (FC 1.12), and *Wdr5* (FC -1.3). In MD-WD testes, we identified 187 down-regulated, 98 up-regulated genes (Figure 5F) associated with an up-regulation of RNA splicing (GO:0008380) and RNA binding (GO:0003723), and a down-regulation of DNA damage (GO:0006974), protein transport (GO:0030154), cell differentiation (GO:0015031) and RNA binding (GO:0003723) pathways (Figure 5G). In contrast to WD testes, we observed an integrated gene network based around the expression of *Snrpg* (FC -1.27), *Top1* (FC 1.38) and *Hnrnph1* (FC -1.3) (Figure 5H) in MD-WD testes. We also observed networks linked to *H3f3a* (FC -1.14), *Mdm2* (FC 1.24) and *Pcgf2* (FC -1.52) in MD-WD testes. As we identified no differentially expressed genes in LPD (Figure 5A) or MD-LPD (Figure 5B) testes, we conducted gene ontology analysis of genes displaying a fold change <-1 or >1 and a P <0.05. In LPD testes we identified 910 genes (490 down-regulated, 420 up-regulated) associated with an upregulation of metabolic processes and a down regulation of RNA binding and transcription regulation. In MD-LPD testes, we identified 3425 (2095 down-regulated, 1330 up-regulated) genes associated with an upregulation of RNA splicing (GO:0008380) and a down regulation of extracellular structure organization (GO:0043062), T cell activation (GO:0046631) and MAPK regulation (GO:0043410).

**Figure 5.**
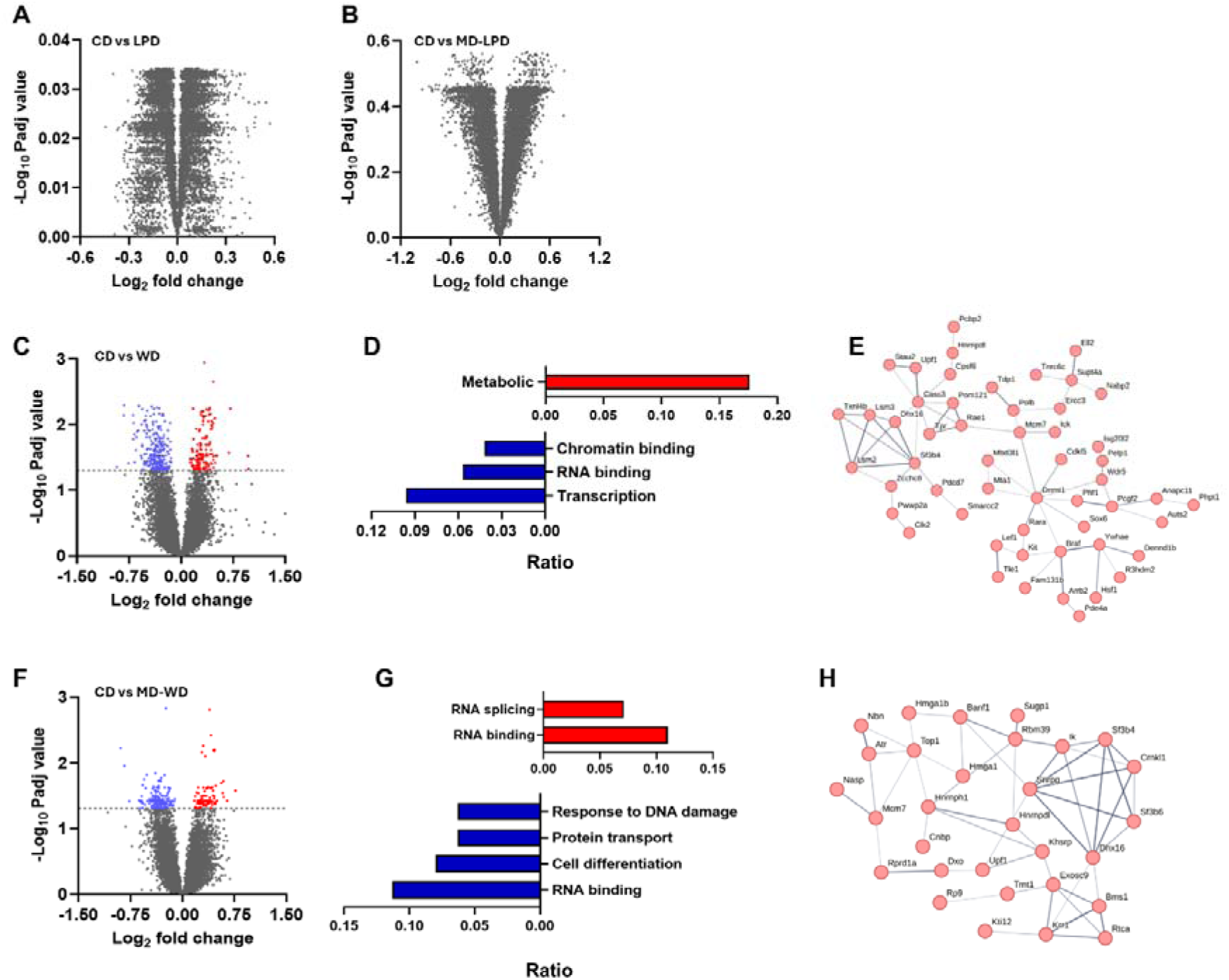
Impact of diet on testicular gene expression. Volcano plots comparing differential gene expression between males fed a control diet (CD) and males fed (A) a low protein diet (LPD), (B) methyl donor-supplemented LPD (MD-LPD) or (C) Western diet (WD). (D) Pathway and (E) network analysis of differentially expressed genes between CD and WD fed males. (F) Volcano plots comparing differential gene expression between males fed a control diet (CD) and males fed a methyl donor-supplemented WD (MD-WD). (G) Pathway and (H) network analysis of differentially expressed genes between CD and MD-WD fed males. N = 8 males in each group.

### Paternal low-protein diet alters early placental morphology and metabolism

Recently, we have shown that while our LPD, MD-LPD, WD and MD-WD diets do not detrimentally affect male fertility, they do have a significant impact on post-fertilisation embryonic development (Morgan et al., 2024). To establish whether sub-optimal paternal diet influenced early placental development, whole conceptus morphology was analysed at embryonic (E) day 8.5. We observed no difference in the mean number of implantation sites between groups (Figure 6A). However, morphological assessment of whole embryo implantation sites (Figure 6B) revealed a significant reduction of invasion depth in LPD embryos compared to MD-LPD embryos (Figure 6C). Furthermore, LPD embryos demonstrated a trend towards a smaller EPC area (Figure 6D) and an increased angle of misalignment from the central maternal channel (Figure 6E) when compared to CD embryos (P <0.1). To explore the impact of paternal diet on early placental dynamics further, individual ectoplacental cones (EPCs) were isolated and cultured for either 24 or 48 hours (Figure 6F). There was no difference in mean trophoblast outgrowth (Figure 6G) or central EPC area (Figure 5H) in either the 24- or 48-hour explants.

**Figure 6.**
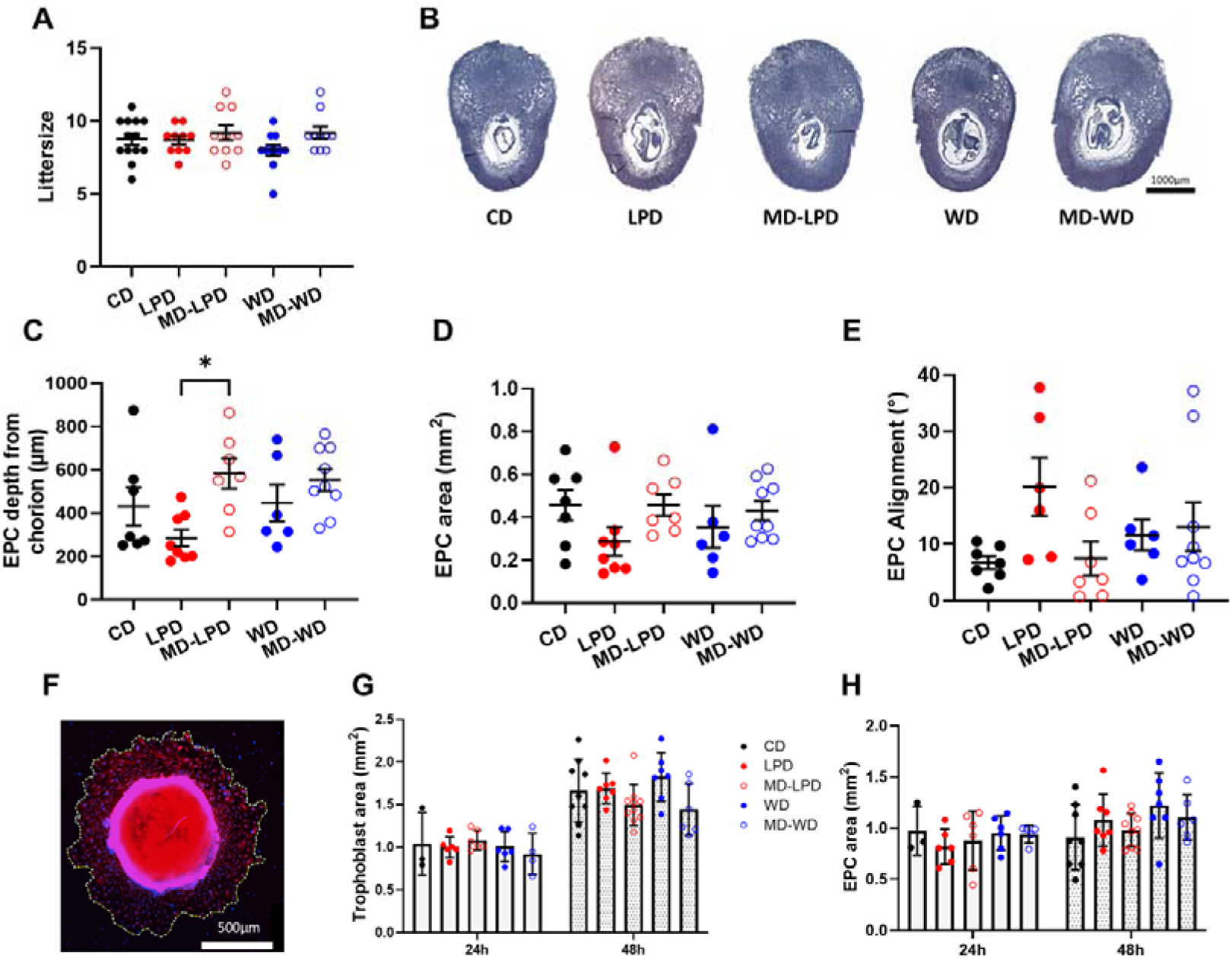
Impact of paternal diet on early (E8.5) placental development. (A) Litter size at E8.5 from males fed either a control diet (CD), low protein diet (LPD), methyl donor-supplemented LPD (MD-LPD), Western diet (WD) or methyl-donor supplemented WD (MD-WD) prior to mating. (B) Representative images of H&E stained whole E8.5 conceptuses. Ectoplacental cone (EPC) (C) invasion dept, (D) area and (I) alignment. (F) Representative EPC outgrowth after 48 hours in culture, stained for alpha-tubulin. (G) Trophoblast and (H) EPC area at 24 and 48 hour timepoints. N = 10-13 litters in A, derived from a minimum of 8 separate stud males per group, and 4-9 conceptuses in C-H, each from a separate litter and stud male. Data were analysed using either a one-way ANOVA or Kruskal–Wallis test with Holm-Sidak or Dunn’s post hoc tests for multiple comparison respectively. * P <0.05.

### Sup-optimal paternal diet alters fetal weights in late gestation in a sex-specific manner

Next, we examined the impact of paternal diets on late gestation (E17.5) fetal and placental development. There was no difference in mean fetal weight between groups. However, in both LPD and MD-LPD groups, a significantly higher proportion of fetuses displayed a weight below the 10^th^ centile when compared to CD fetuses (15% and 19% respectively, P <0.05, Figure 7A). No differences in fetal weight distribution were observed in either WD or MD-WD groups (Figure 7A). When fetal weights were separated by fetal sex, LPD, MD-LPD and MD-WD female fetusess were significantly lighter than CD females (P <0.05; Figure 7B). However, there were no differences in mean fetal weight between any of the male groups (Figure 7C). When assessing the influence of fetal sex on weight, we observed that males were typically heavier than females, with the difference being significant for CD, LPD, WD and MD-WD males when compared to their respective females (P <0.05, Figure 7D).

**Figure 7.**
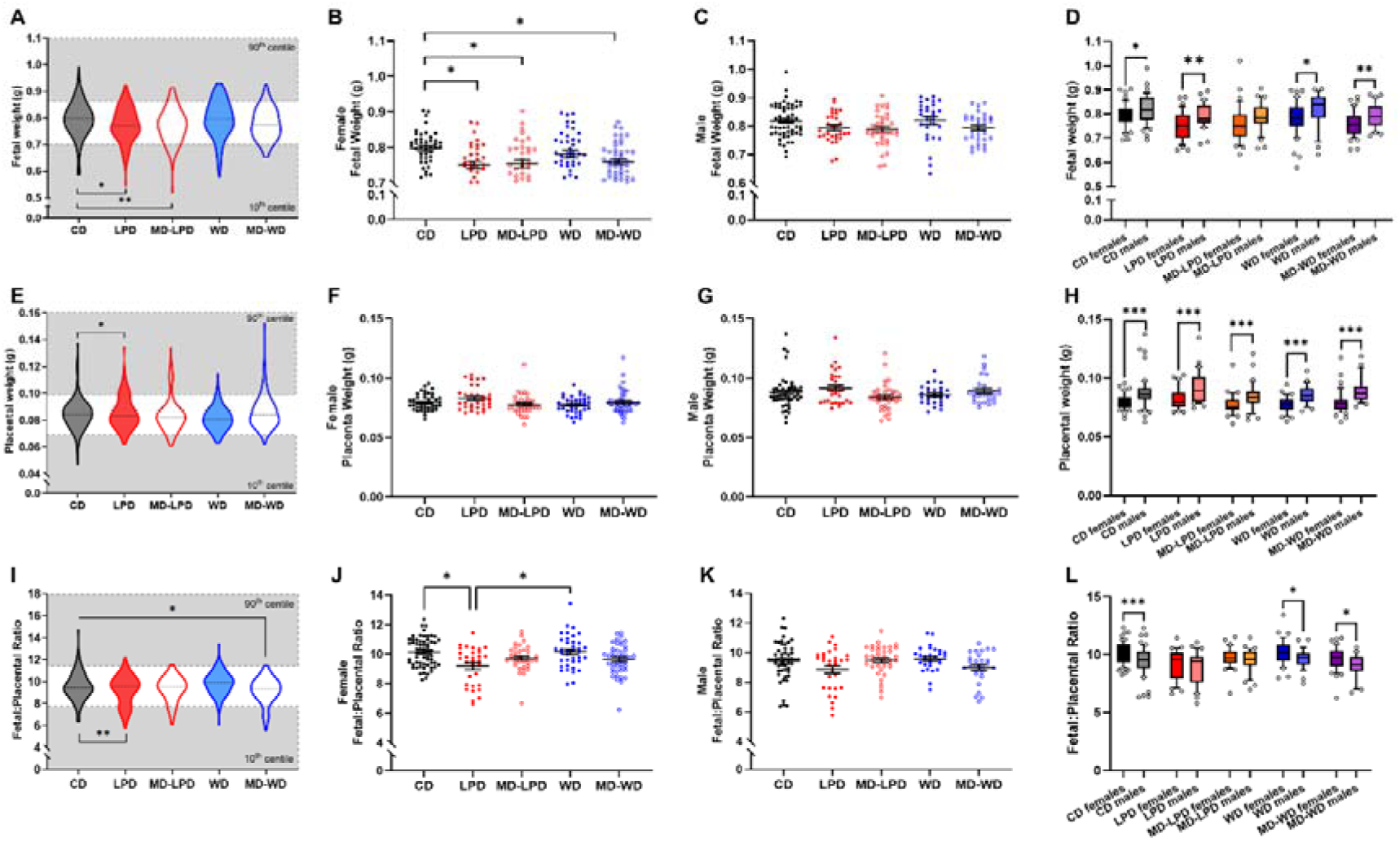
Impact of paternal diet on fetal and placental weight in late (E17.5) gestation. (A) Late gestation fetal weight and distributions above the 90^th^ centile and below the 10^th^ centile of control diet (CD) fetal weights from males fed either a CD, low protein diet (LPD), methyl donor-supplemented LPD (MD-LPD), Western diet (WD) or methyl donor-supplemented WD (MD-WD) prior to mating. Late gestation (B) female, (C) male and (D) intra-diet comparison of fetal weight. (E) Late gestation placental weight and distributions above the 90^th^ centile and below the 10^th^ centile of CD placental weights. Late gestation (F) female, (G) male and (H) intra-diet comparison of placental weight. (I) Late gestation fetal:placental ratio and distributions above the 90^th^ centile and below the 10^th^ centile of CD fetal:placental ratio. Late gestation (G) female, (H) male and (I) intra-diet comparison of fetal:placental weight ratio. N = 10-13 litters, derived from a minimum of 8 separate stud males per group. Data are presented as mean ± SEM or as box plots with the mean and individual data points outside of the 10-90^th^ percentile. Data were analysed using a generalised linear mixed model analysis with paternal origin of litter and duration on respective diet incorporated as random effects (panels A, B, C, E, F, G, I, J and K) or t-test (panels D, H and L) following assessment for normality using a Shapiro-Wilk test. * P <0.05, ** P <0.01, ***P <0.001.

Analysis of mean placental weight showed no difference between groups. However, placentas from LPD fed males showed a higher proportion above the 90^th^ centile for weight when compared to CD males (Figure 7E, P <0.05). Analysis of placental weight by fetal sex showed no difference for either females (Figure 7F), or males (Figure 7G) in any groups. However, in line with fetal weight, males had heavier placentas than females in all groups (P <0.001, Figure 7H).

Finally, while there was no difference in mean fetal:placental ratio across all groups, a significantly higher proportion of LPD fetuses displayed a ratio below the 10^th^ centile, and MD-WD fetuses showed a reduction in the number above the 90^th^ centile (Figure 7I; P <0.05), when compared to CD fetuses. Analysis of fetal:placental ratio revealed a significantly lower ratio in LPD females when compared to CD and WD females (P <0.05, Figure 7J). In contrast, no differences between any of the male groups was observed (Figure 7K). When comparing females to males, we observed a lower fetal:placental ratio in CD, WD and MD-WD males (P <0.05, Figure 7L) when compared to their respective females. However, no differences were observed between LPD or MD-LPD males and females (Figure 7L).

### Paternal sub-optimal diet has minimal impact on placental morphology

As fetal placental ratio appeared altered in response to paternal diet, we next assessed placental morphology, measuring the relative areas of the decidua, junctional zone, labyrinth zone and chorionic plate (Figure 8A). There was no difference in mean cross sectional area (Figure 8B) or in the relative proportion of each placental compartment (Figure 8C) between diets. Additionally, we undertook a stereological analysis of placenta morphology (Figure 8D-F) observing no difference in mean placental volume (Figure 8G), labyrinth zone volume (Figure 8H), junctional zone volume (Figure 8I), maternal blood volume (Figure 8J), maternal blood surface area (Figure 8K) or fetal capillary surface area (Figure 8L).

**Figure 8.**
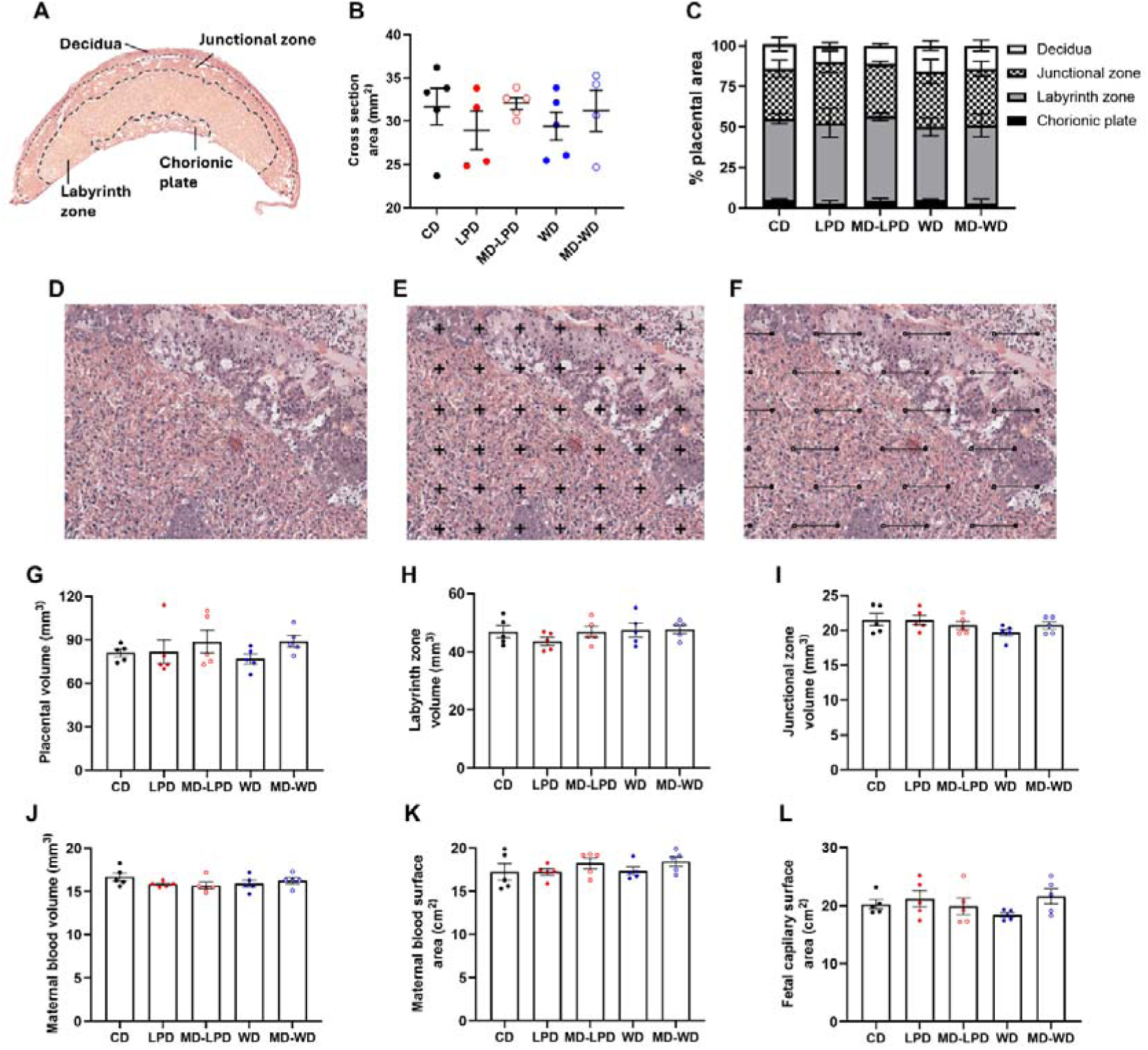
Analysis of late gestation (E17.5) placental morphology in response to paternal diet. (A) Representative image of a cross section through the late gestation (E17.5) placenta highlighting the decidua, junctional zone, labyrinth zone and chorionic plate regions. (B) Cross section area and (C) relative regional composition of placentas derived from males fed either a control diet (CD), low protein diet (LPD), methyl donor-supplemented LPD (MD-LPD), Western diet (WD) or methyl-donor supplemented WD (MD-WD) prior to mating. (D) Representative images of a E17.5 placenta at 20X magnification prior to the application of (E) estimation of compartment global volumes using Cavalieri’s Principle with a grid comprising a test system of points and (F) stereological method for estimating the surface density of maternal blood vessels and fetal capillaries using a grid comprising isotropic lines. E17.5 placental (G) volume, (H) labyrinth zone volume, (I) junctional zone volume, (J) maternal blood space volume, (K) maternal blood surface area and (L) fetal capillary surface area. N = 5 placentas per group, each from a separate litters and stud males. Data are presented as mean ± SEM and were analysed using a one-way ANOVA (panels B, G-L) or Kruskal–Wallis test (panel B) with post-hoc correction where appropriate.

### Paternal sub-optimal diets influence sex-specific placental transcriptome

Finally, we performed a transcriptomic analysis of the late gestation placenta by RNA-Seq using 4 male and 4 female placentas from each dietary group. Initial analysis, separating the results by diet and sex, identified 0, 102, 34 and 77 differentially expressed genes (Padj <0.05) between LPD, MD-LPD, WD, MD-WD and CD male placentas respectively (Figure 9A). Pathway analysis of the 102 differentially expressed genes in MD-LPD male placentas identified pathways involved in metabolism (Type 1 Diabetes mellitus), immune system regulation (antigen processing, complement and coagulation cascades), cellular adhesion, signalling and senescence (Figure 9B). No significant pathway enrichment was found for the differentially expressed genes in WD or MD-WD male placentas when compared to CD males. Similarly, when comparing female placentas, we observed 0, 3, 97 and 52 differentially expressed genes (Padj <0.05) when comparing LPD, MD-LPD, WD and MD-WD with CD placentas respectively (Figure 9C). Pathway analysis of the 97 differentially expressed genes in WD female placentas identified pathways involved in lipid metabolism and signalling (RAS, GnRH) pathways (Figure 9D). No significant pathways were identified for either MD-LPD or MD-WD female placenta differentially expressed genes. These data suggest that paternal diet may be influencing placental gene expression in a sex-dependent manner, with more differentially expressed genes observed within the male MD-LPD and MD-WD groups, while the WD and MD-WD females shower a greater change in gene expression profiles. Principal component analysis of expression between CD males and females suggested the existence of a sex-specific profile (Figure 9E). Additional analysis identified 301 differentially expressed placental genes between CD males and females (Padj <0.05, Figure 9F) (See Supplemental Data S1 for full list). Of these 301 genes, 43 were upregulated in males and 258 were upregulated in females. Gene ontology analysis identified pathways involved in vascular development, developmental processes and signal transduction (Figure 9G). Interestingly, comparison of these same 301 genes in LPD, MD-LPD, WD and MD-WD placentas showed no such sexual dimorphism (Figure 9H-K). In total, we identified 13, 0, 14 and 15 genes to be differentially expressed between males and females in the LPD, MD-LPD, WD and MD-WD groups respectively (Figure 9L-O). In total, we identified just 9 differentially expressed genes, and one unprocessed pseudogene, present in all 5 dietary groups (Figure 9P). Of the 9 conserved differentially expressed mRNAs, 5 were located on the X chromosome (*Eif2s3x, Taf1, Otg, Xist, Kdm5c*), while 4 were on the Y chromosome (*Kdm5d, Uty, Ddx3y, Eif2s3y*) (Figure 9Q). Separate to these, LPD placentas displayed sexually dimorphic gene expression on chromosomes 5 and 13, WD placentas had sexually dimorphic gene expression on chromosomes 3, 5 and 15, and MD-WD demonstrated sexually dimorphic gene expression on chromosomes 3, 5, 6 and 14 (see Supplementary Data S1).

**Figure 9.**
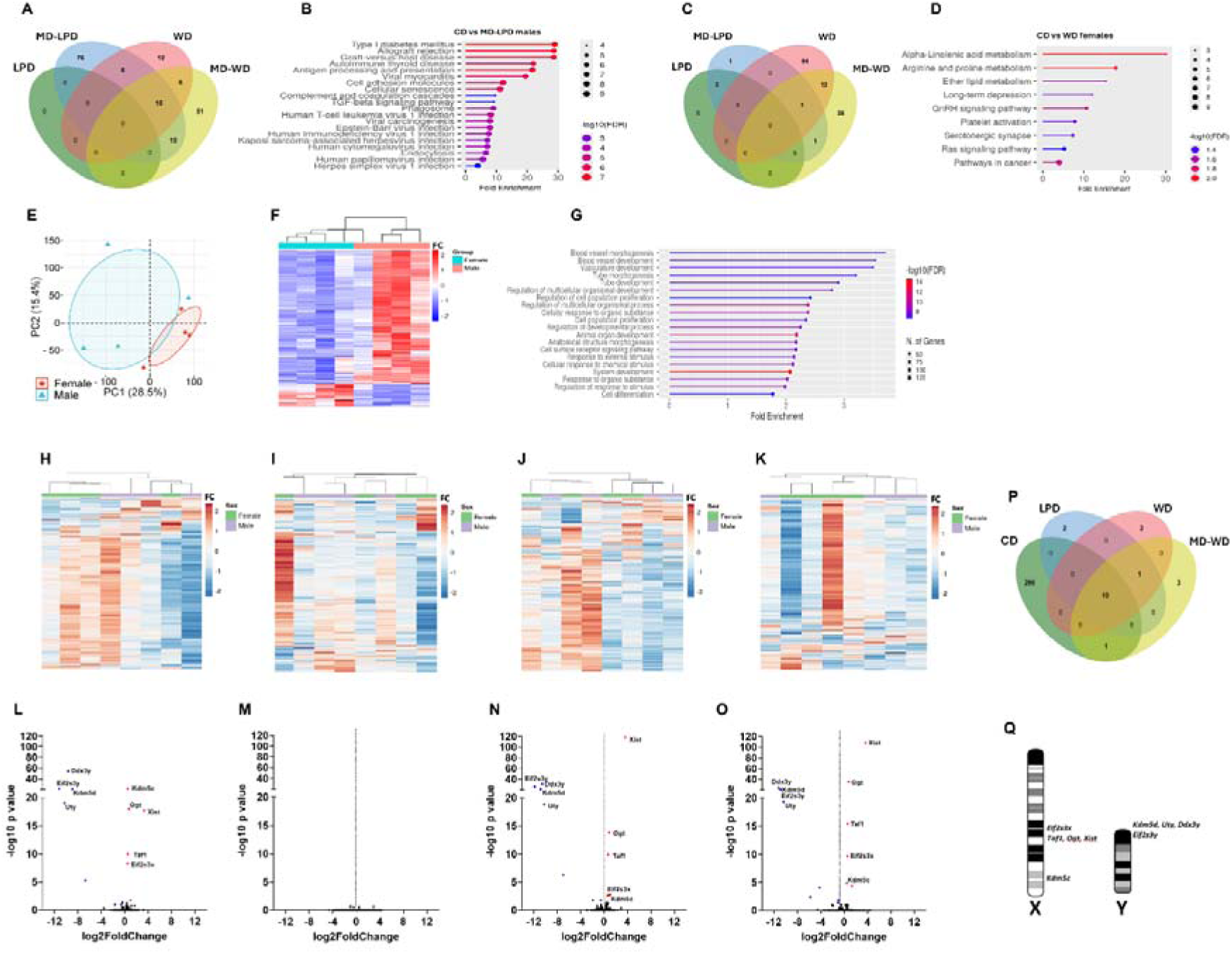
Late gestation (E17.5) placental gene expression is altered in a diet and sex-specific manner. (A) Differential gene expression and (B) pathway analysis from male placentas derived from males fed either a control diet (CD), low protein diet (LPD), methyl donor-supplemented LPD (MD-LPD), Western diet (WD) or methyl-donor supplemented WD (MD-WD) prior to mating. (C) Differential gene expression and (D) pathway analysis from female placentas. (E) Principal component analysis comparing CD male and female placental gene expression profiles. (F) Heat map of the 301 sexually dimorphic genes between male and female CD placentas and (G) the pathway analysis. Heatmaps showing the relative level of expression and clustering of the same 301 genes in (H) LPD, (I) MD-LPD, (J) WD and (K) MD-WD male and female placentas. Volcano plots highlighting the significantly (pAdj <0.05) upregulated (red) and downregulated (blue) differentially expressed genes between male and female (L) LPD, (M) MD-LPD, (N) WD and (O) MD-WD placentas. (P) Overlap of sexually dimorphic genes between male and female CD, LPD, WD and MD-WD placentas. (Q) Chromosomal locations of common X and Y chromosome sexually dimorphic genes between CD, LPD, WD and MD-WD placentas. N = 4 male and 4 female placentas, each from separate litters and stud males.

## Discussion

Consumption of an unbalanced or unhealthy diet is a major risk factor for a range of diseases such as obesity, diabetes and cardiovascular disease. Poor parental diet prior to conception also impacts on patterns of post-fertilisation development, shaping fetal and placental dynamics and increases the risk for a range of non-communicable disorders in adult life (Ozanne et al., 2004). Under the current study, we explored the impact of both paternal under-(LPD) and over-nutrition (WD) on male reproductive health, fetal growth and placental development. We also investigated the impact of supplementing these diets with a range of methyl-donors and carriers to define if their use could negate any detrimental influences imposed by the diets. We observed that males consuming a high fat/sugar diet (WD and MD-WD), displayed a series of physiological changes in their metabolic health, gut bacterial profiles and testicular morphology and gene expression. While both over- and under-nutritional regimens had no impact on fundamental male fertility, we observed significant changes in fetal and placental weights in late gestation. Furthermore, we observed a significant loss in sexual dimorphism in the late gestation placenta. While sex-specific differences in fetal and adult offspring are reported widely in response to both maternal and paternal diet (Dearden et al., 2018, Rosenfeld, 2015), our data provide additional insight into how sex-specific paternal programming may be mediated during development.

While we observed minimal impact of the LPD and MD-LPD on male metabolic health, mice fed the WD and MD-WD displayed significant changes in adiposity and hepatic cholesterol and FFAs. Excessive accumulation of cholesterol and FFAs within the liver are central hallmarks of metabolic conditions such NAFLD and MASLD. While elevated hepatic FFAs have been linked to insulin resistance and elevated levels of proinflammatory mediators, we observed no change in circulating glucose, insulin or Tnf levels. This is in line with studies showing that nearly half of patients with NAFLD/MASLD also do not present with insulin resistance (Singh et al., 2015), highlighting it heterogenous nature. Changes in metabolic status have also been linked to gut dysbiosis. We observed differences in the abundance of Defferibacteres, Proteobacteria and TM7 in fecal samples from WD and MD-WD males. Similar increases in Deferribacteres have been observed in mice maintained on high fat diets (Liu et al., 2019, Serino et al., 2012), while the abundance of TM7 has been linked with elevated BMI, fat mass and inflammatory status in humans (Gomes et al., 2020). Separately, negative links between TM7 abundance and body weight and adiposity in rodents have been identified (de la Garza et al., 2022, Hu et al., 2022). While our study indicates significant shifts in male metabolic homeostasis, further studies are needed to define how metabolic, inflammatory and microbiota status interact and their impact on male reproductive health. For example, bacterial metabolites such as 5-hydroxytryptamine (5-HT) have been shown to promote sperm hyperactivation and the acrosome reaction in mice (Sugiyama et al., 2019), hamsters (Fujinoki, 2011) and men (Omote et al., 2023). Additionally, many polyunsaturated fatty acids (PUFAs) produced by the gut microbiota have significant role in stabilising the sperm plasma membrane (Lv et al., 2024). Finally, additional microbiota derived metabolites such as androgens (Collden et al., 2019) and hormones including GLP-1 and peptide YY (PYY) (Cannarella et al., 2021) have been associated with changes in sperm production. Interestingly, studies have also shown that fecal transplants can be used to treat infertility and improve sperm quality in mice (Hao et al., 2022, Zhang et al., 2021). Therefore, additional studies, with the inclusion of a methyl donor CD (MD-CD), are required to understand better the connections between dietary-induced gut dysbiosis and male reproductive fitness.

Within the testis, LPD and WD males displayed a significant reduction in the number of undifferentiated spermatogonial stem (Plzf+) cells when compared to CD males, while no difference was observed in MD-LPD and MD-WD males. While the loss of Plzf is not detrimental for spermatogenesis, Plzf ensures spermatogonial stem remain in a state of undifferentiation, interacting with other transcription factors such as SRY-box TF 3 (SOX3) and Spalt like TF 4 (SALL4) to establish the chromatin and transcription landscape in spermatogonial stem cells (Yi et al., 2025). However, the regulation of Plzf by methyl donors is currently undetermined. Separately, WD and MD-WD males displayed an increased rate of tubule abnormalities when compared to CD males. In rodents, exposure to a high fat diet has been shown to induce tubule vacuolization atrophy (Ghosh and Mukherjee, 2018) as well as disrupting the essential adhesion between spermatogenic and Sertoli cells, resulting in Sertoli cell atrophy (Liu et al., 2014). Sertoli cells are essential for supporting spermatogenesis and their loss is associated with male infertility. Sertoli cells also produce inhibin β and its serum levels correlate positively with testicular size and sperm production rates (Luisi et al., 2005).

Inhibin production is also tightly linked to the secretion of FSH from the pituitary, providing a negative feedback signal to regulate spermatogenesis. As we observed a significant elevation in serum inhibin β-A in WD males, but no change in Sertoli (Sox9^+^) cell number or fundamental male fertility, the increase in inhibin β might indicate a dysfunction in normal hypothalamic-pituitary-testis axis regulation. However, further studies are needed to confirm this.

Analysis of testicular gene expression revealed minimal changes in response to LPD or MD-LPD. In contrast, testes from WD and MD-WD-fed males displayed an upregulation of genes involved in fatty acid metabolism, mRNA processing, Wnt signalling and protein transport processes, and a downregulation of transcription related genes. The increase in genes associated with fatty acid metabolism (*Acadsb*, *Crat, Pla2g5*, *Daglb, Mgll*) and glycolytic components (*Pdha1, Hk2*) link with the changes observed in metabolic status in these males. Furthermore, we observed up regulation of *Paf1* and *Ctr9,* both components of the Paf1C complex. PAF1C associates with the promoter and coding sequences of active genes (Pokholok et al., 2002) and interacts with a series of transcriptional elongation factors (Pokholok et al., 2002, Squazzo et al., 2002). Studies have identified Paf1C as crucial for the acquisition of transcription-associated histone modifications such as H3 methylation at K4, K36, and K79 (Cao et al., 2015, Mbogning et al., 2013). Interestingly, we identified an increase in the expression of the histone variants *H3f3a* and *H3f3b*. Unlike most other canonical histones, H3.3 is not completely removed from the chromatin during spermatogenesis and is enriched at genomic regions important for zygotic development in mice and men (Brykczynska et al., 2010, Hammoud et al., 2011). Studies have shown that disruption of H3K4me2 marks in mouse sperm have significant consequences for offspring health over multiple generations (Siklenka et al., 2015) and levels of sperm H3K4me2 may serve as a metabolic sensor linking paternal diet with offspring metabolic ill-health predisposition (Pepin et al., 2022). The downregulation of transcription regulating genes such as *Dnmt1, Mta1, Phf1, Pwwp2a, Wdr5, Hsf1, Nfrkb. Pelp1, Prox1* and *Rara* suggest paternal HFD might influence differential patterns of testicular transcriptional regulation, which could affect sperm composition and post-fertilisation development (Cui et al., 2025). Finally, we observed a down regulation of multiple RNA binding genes in both WD and MD-WD testes. Recently, the significance of RNA-binding proteins as critical regulators of spermatogenesis and sperm function has been recognised (Li et al., 2024). As many RNA-binding proteins play central roles in orchestrating intricate transcriptional and post-transcriptional regulatory networks, their function as essential regulators of testicular germ cell development is becoming defined (Gao et al., 2026).

Together, these findings suggest that sub-optimal male diets have the potential to alter sperm modifications, which are not simply corrected by methyl-donor supplementation. To determine whether these paternal diet-driven changes to male reproductive physiology impacted on post-fertilisation dynamics, we analysed early (E8.5) and late (E17.5) fetal and placental development. Analysis of E8.5 placental invasion revealed poor paternal diet had minimal impact on overall morphology or invasion depth *in vivo*. However, we observed significant sex-specific influences of paternal diet on late gestation fetal and placental growth. During fetal development, differences in patterns of male and female growth have been described from as early as the first trimester with males growing at a faster rate (Broere-Brown et al., 2016). We identified over 300 sexually dimorphic, differentially expressed genes between CD males and females, with the majority being upregulated in females. Gene ontology analysis identified vascular, immunological, extracellular matrix organisation and cell-signalling pathways as being differentially expressed, mirroring observations from male and female term placentas in humans (Buckberry et al., 2014). While female human term placentas have increased expression of multiple immune regulation genes when compared to males (Sood et al., 2006), Y-chromosome genes such as *Ddx3y, Uty,* and *Kdm5d,* encode epitopes that contribute to minor histocompatibility antigens present throughout the placenta (Cvitic et al., 2013). Studies have also shown that cultured trophoblast tissues isolated from human healthy male placentas secrete more TNF and less IL-10 in response to lipopolysaccharide than female tissues (Yang et al., 2021). In contrast, genes such as *Tgfb1,* which was upregulated in CD females, are critical in regulating fetal-maternal immune tolerance (Yang et al., 2021). Furthermore, studies have shown female placentas to be more responsive to shifts in maternal inflammation and dietary status than males (Osei-Kumah et al., 2011, Sedlmeier et al., 2014). Similar to our data, studies have also reported upregulation of receptor-ligand signalling in human female placentas including several collagens, laminins and integrins, suggestive of an enhanced invasion and placentation potential (Braun et al., 2021). Finally, we observed up regulation of genes involved in steroid synthesis and endocrine regulation in female placentas. Placental endocrine function operates to regulate both maternal physiological response and fetal development (Stern et al., 2021). In contrast to our CD males and females, all other groups displayed a dramatic loss of sexual dimorphism. While the underlying mechanism(s) for our observed loss of sexual-dimorphism are unknown, perturbed paternal sperm epigenetic status could modulate early embryo X-chromosome inactivation and/or X-chromosome dosage in females, resulting in differential profiles of autosomal gene expression (Gonzalez et al., 2018, Petropoulos et al., 2016).

Collectively, we observe that paternal LPD or WD, with or without supplementation, had no impact on overall male fertility, but have influences on male reproductive physiology. While paternal LPD induced minor changes in early placental morphology and metabolism, these effects were small, and the consequences of these changes remain to be defined. In late gestation, paternal diet affected fetal development in a sex- and diet-specific manner. While we observed significant sexual dimorphism in gene expression patterns between CD males and females, these differences were largely removed in response to our experimental diets. Such loss of sexual- dimorphism could provide one process through which poor paternal diet programs offspring ill-health in adulthood. However, the precise mechanisms through which this occurs remain to be determined.

## Materials and Methods

### Mice diet regime and matings

All animal procedures were conducted in accordance with the UK Home Office Animals (Scientific Procedures) Act 1986 and carried out under Project License PP8899264 with local ethical approval at University of Nottingham. C57BL/6J mice (Charles River, UK) were housed in controlled 12/12-hour light/dark conditions with a constant temperature (21° C ± 3°C) and *ad libitum* access to water. Virgin 8-week old males (n = 8 per treatment group) were fed either control diet (CD; 18% casein, 21% sucrose, 0% milk fat, 0% cholesterol), isocaloric low protein diet (LPD; 9% casein, 24% sucrose, 0% milk fat, 0% cholesterol), ‘Western’ diet (WD; 19% casein, 34% sucrose, 20% milk fat, 0.15% cholesterol) or LPD or WD supplemented with methyl-donors and carriers (5 g/kg diet choline chloride, 15 g/kg diet betaine, 7.5 g/kg diet methionine, 15 mg/kg diet folic acid, 1.5 mg/kg diet vitamin B12; termed MD-LPD or MD-WD respectively), for up to 24 weeks. Exact dietary formulations are outlined in Supplementary Table S1. Virgin female C57BL/6J mice were maintained on standard rodent chow (rat/mouse No.1 maintenance diet, Special Diet Services) throughout study the study. Females were mated at 9-weeks old (+/-7 days) with stud males who had been fed their respective diet for a minimum of 8 weeks. Successful mating was confirmed by the presence of a copulation plug and denoted as embryonic day (E)0.5. Dams were euthanized via cervical dislocation on either E8.5, for the collection of ectoplacental cones (EPCs), or E17.5 for fetal and placental tissues. Stud males were culled by cervical dislocation for the collection and storage of tissues. Blood was collected via heart puncture, allowed to clot on ice and centrifuged at 10,000 × g, 4 °C for 10 minutes before storage of the serum at −80 °C. Heart, kidney, liver, testis, and caudal epididymis were dissected, weighed and either snap frozen prior to storage at −80 °C, or fixed overnight in 10 % neutral buffered formalin (Sigma, UK) at 4 °C before processing and embedding in paraffin wax. Fecal pellets were collected from the descending colon using sterile forceps and placed in 2 ml DNase/RNase-free collection tubes, snap frozen and stored at -80 °C. A second batch of males (n = 6) were fed using the same experimental protocol and utilized for the analysis of fat pad mass.

### Stud male metabolic status assessment

Stud male non-fasting serum glucose was measured using the Glucose Colorimetric Detection Kit (EIAGLUC, Thermo Fisher Scientific, UK), non-fasting insulin was measured using a Rat/Mouse Insulin ELISA Kit (EZRMI-13K; Millipore, UK), Tnf was measured using the TNF alpha SimpleStep ELISA® Kit (ab208348, Abcam®), inhibin β-A chain was measured using the FineTest® Mouse Inhba ELISA Kit (EM0273, Wuhan Fine Biotech Co. China), non-fasting liver tissue total cholesterol was measured using the Cholesterol Quantification Kit (MAK043, Sigma-Aldrich, UK), non-fasting hepatic free fatty acids (FFA) were measured using the Free Fatty Acid Quantitation Kit (MAK044, Sigma-Aldrich, UK) and non-fasting hepatic triglycerides were measured using the Triglyceride Quantification Kit (MAK266, Sigma-Aldrich, UK), all in accordance with the manufacturers’ instructions.

### Gut microbiota sequencing

DNA was extracted from stool pellets using the QIAamp DNA Stool Mini Kit (Qiagen, UK) following the manufacturer’s instructions and stored at -80 °C. 16S rDNA sequencing (V3-V4 region) was conducted as described previously (Morgan et al., 2022). The extracted stool DNA samples (n = 6 per diet group) were sequenced using the Illumina MiSeq Reagent kit v3 on the Illumina MiSeq platform (Illumina), in accordance with Illumina 16S Metagenomic Sequencing Library Preparation protocol. Briefly, 16S rRNA amplicons were generated using the forward 5’ (TCGTCGGCAGCGTCAGATGTGTATAAGAGACAGCCTACGGGNGGCWGCAG) and Reverse 5’ (GTCTCGTGGGCTCGGAGATGTGTATAAGAGACAGGACTACHVGGGTATCTAATC C) primers, flanked by Illumina adapter-overhang sequences. Illumina dual index barcodes (Illumina XT Index Kit v2, Set A: FC-131-2001) were attached to each amplicon. PCR clean-up was conducted using AMpure XP beads (Beckman; A63882). Library fragment-length distributions were analysed using the Agilent TapeStation 4200 and the Agilent D1000 ScreenTape Assay (Agilent; 5067-5582 and 5067-5583). The generated libraries were pooled in equimolar amounts and the pool was size selected using the Blue Pippin (Sage Science) and a 1.5 % Pippin Gel Cassette (Sage Science; BDF2010). The samples were then run over a shared 300 paired end (PE) MiSeq run to deliver about 60-80,000 PE reads per sample. The sequencing run additionally had a 20 % PhiX library spike-in as an internal quality control. Raw reads were processed by Qiime2 pipeline and trimmed. Greengenes version 13.8 was used in the classification (Caporaso et al. 2010).

### Testicular histology

Parafin embedded testis tissue (n = 8 per group) was sectioned at 5 μm using a Leitz 1512 rotary microtome (Leica). For analysis of seminiferous tubule morphology, sections were processed and stained with Haematoxylin-Eosin prior to imaging using a Leica DMRB microscope with an Oasis glide scanner. Image analysis was carried out using Fiji. For each testis sample, an average of 50 seminiferous tubule cross-sections were analysed.

For the determination of seminiferous tubule cell types, additional testis sections were dewaxed, rehydrated into PBS, and equilibrated in 100 mM citrate buffer (pH 6.0) at room temperature (RT) prior to being microwaved in 0.1M tri-sodium citrate buffer (pH 6.0) for 5 minutes. The sections were incubated in blocking solution (5 % bovine serum albumin, 20 % normal goat serum in 0.1 % Triton X-100 in PBS; Sigma, UK) for 1 hour before incubating with anti-Ddx4 antibody (to identify germ cells; ab27591, 1:100; Abcam, UK) or Rabbit anti-Sox9 Antibody (to identify Sertoli cells: ab5535, 1:500; Chemicon, UK) in blocking solution, and incubated at 4° C overnight. Slides were washed (three times in 0.1% Triton X-100 in PBS; PBST) prior to incubation in Alexa Fluor 568 (A-21124, 1:200; Invitrogen, UK) in blocking solution for the detection of Ddx4. For the detection of Sox9, slides were incubated with goat anti-rabbit biotinylated antibody (E0432, 1:200; DakoCytomation, UK) for 1 hour in the dark in blocking solution prior to Alexa Fluor 488 streptavidin conjugate, washed in PBST prior to incubation with Alexa Fluor 488 (S-32354, 1:200; Invitrogen, UK) in blocking solution for 30 minutes at room temperature. For the detection of Plzf, sections were incubated overnight with anti-Plzf antibody (to identify spermatogonial stem cells: ab189849, 1:3000; Abcam, UK) at 4 °C. Sections were washed in PBST prior to incubation with Alexa Fluor 568 (ab175471, 1:200; Abcam) in blocking solution for 1 hour at room temperature. Negative controls (absence of primary or secondary antibodies) were also conducted to confirm staining specificity (see Supplemental Figure 1). Slides were washed (PBST) before mounting with Antifade Mounting Media with DAPI (Vectashield, UK). Sections were imaged using a Nikon Eclipse 90i fluorescent microscope with a mercury-fiber illuminator Nikon Intensilight C-HGFI and a Hamamatsu ORCA-ER Digital camera (C4742-80) at 10 and 20 x magnification. Images were analysed using Volocity (Quorum Technologies Inc) with an average of 50 seminiferous tubules counted per male.

### Testicular micro-array

Total RNA was extracted from frozen testicular tissues using the RNeasy Plus Mini Kit (Qiagen; UK) following the manufacturer’s instructions. Testicular RNA was diluted to 100 ng / μl using RNAse-free water (Qiagen, UK) prior to RNA integrity assessment on a Bioanalyzer 2100 platform (Agilent, UK). Samples with an RIN >7 were used for analysis. The GeneChip™ WT PLUS Reagent Kit (ThermoFisher Scientific, UK) was used to prepare RNA samples for whole transcriptome expression analysis with GeneChip™ Whole Transcript (WT) Expression Arrays, as per the manufacturer’s instructions. First-strand cDNA was synthesized from the extracted RNA using Reverse Transcriptase, followed by a second-strand cDNA synthesis using DNA Polymerase and RNAse H. Complementary RNA (cRNA) was synthesized by *in vitro* transcription of the second-strand cDNA using T7 RNA polymerase and purified. The purified ss-cDNA was enzymatically fragmented and labelled by terminal deoxynucleotidyl transferase (TdT) using the provided DNA Labelling Reagent following the manufacturer’s instructions prior to hybridisation to onto the Clariom™ S Assay Mouse GeneChip™ (ThermoFisher Scientific, UK). Data analysis was performed using the Partek Genomics Suite 6.6 analysis software. Comparative analysis between the five diet groups was performed using standard One-way ANOVAs. Dysregulated transcripts were classified based on False Discovery Rate (FDR) = 0.05, fold change < +/-1.1, and p ≤ 0.05. The functional annotation and enrichment analysis web tool WEB-based GEneSeTAnaLysis Toolkit (WebGestalt) (Liao et al., 2019) was used for Gene set enrichment analysis (GSEA). Network analysis was conducted using Network Analyst (https://www.networkanalyst.ca/NetworkAnalyst/home.xhtml). All gene expression data is publicly available via Gene Expression Omnibus (GSE279868).

### *In vivo* EPC invasion

Formalin-fixed, paraffin-embedded E8.5 whole implantation sites were sectioned to the mid-point as indicated by appearance of the maternal channel. Haematoxylin-Eosin staining was conducted on 5 µm sections and whole sites were imaged at 20x magnification using Leica a DMRB microscope with an Oasis glide scanner. Images were imported into Fiji for the assessment of total EPC, trophoblast specific area, depth of invasion (measured from chorion to the furthest trophoblast going in the direction of the maternal channel) and alignment of invasion measured according to (Wilkinson et al., 2021) within the implant site and was conducted by a blinded observer.

### Isolation and culture of EPC explants

The isolation and culture of E8.5 EPCs was conducted as previously described (Watkins et al., 2015). Twenty-four hours prior to dissection, sterile coverslips were coated with BD Matrigel basement membrane matrix (BD Biosciences, Oxford, UK) diluted to 6□mg/ml in RPMI 1640 medium (Life Technologies, UK) and kept at 4° C in four-well plates. On the morning of EPC isolation, the Matrigel-RPMI medium was replaced with 500□μl sterile filtered RPMI-1640 medium containing 2% KnockOut Serum Replacement (Life Technologies, UK), penicillin–streptomycin–glutamine mix (Life Technologies, UK) and 26.2□μmol 2-mercaptoethanol (Sigma–Aldrich, UK) and allowed to equilibrate at 37□° C in 5% CO2 for at least 1□hour. Whole uteri were excised and placed in pre-warmed (37° C) DMEM (Fisher Scientific, UK) supplemented with 10% (v/v) fetal calf serum (Life Technologies, UK). Individual implantation sites were dissected from the uterine tissue and placed in RPMI-1640 medium supplemented with 2% KO-serum replacement (Fisher Scientific, UK), 1% penicillin–streptomycin–glutamine mix (Fisher Scientific, UK) and 26.2µmol 2-mercaptoethanol (Sigma–Aldrich, UK). The EPCs was then isolated from the decidual capsules and separated from the embryo. The isolated EPCs were then placed onto the Matrigel-coated coverslips and cultured individually at 37□° C, 5% CO_2_ for either 24 or 48 hours, with a 50% medium change after 24□hours.

After 24 or 48 hours in culture, EPC outgrowths were fixed in 4% neutral buffered formalin (Sigma–Aldrich, UK) for 15 minutes at room temperature, washed (1x PBS) and stored at 4° C in PBS. Fixed EPCs were permeabilised with 0.1% Triton-X100 (Sigma–Aldrich, UK) and autofluorescence quenched with ammonium chloride (2.6□ mg/ml in PBS; Sigma-Aldrich UK). Non-specific binding was blocked with 2% BSA in PBS containing 0.1% Triton-X100 for 30□minutes at room temperature before overnight incubation at 4□° C with a primary antibody for α-tubulin (1:2,000; Cell Signaling, Danvers, MA, USA) in PBS with 1% BSA and 0.1% Triton-X100. Following washing (PBS, 0.1% Triton-X100), outgrowths were incubated with the appropriate Alexa Fluor conjugated secondary antibody (1:10□,000; Molecular Probes, Life Technologies, Paisley, UK) for 1□hour at room temperature and counterstained with DAPI (Sigma–Aldrich, UK) for 10□minutes before mounting on slides in DPX (Fisher Scientific, Loughborough, UK). All outgrowths were imaged using a Nikon Eclipse 90i fluorescent microscope fitted with a mercury-fiber illuminator, Nikon Intensilight C-HGFI and a Hamamatsu ORCA-ER Digital camera (C4742-80) and analysed using the Volocity Software. The central EPC was defined as the central, single mass within which individual cell nuclei could not be distinguished from those neighbouring them. Proliferative trophoblast nuclei were defined as those located within the proximity of the central EPC as described previously (Watkins et al., 2015).

### Placental stereology

At E17.5, dams were culled for the collection of placental tissues. Fetuses and placentas were weighted prior to fixation in Karnovsky’s fixative (2.5% glutaraldehyde, 2% formaldehyde in 0.1M sodium phosphate buffer) for 16 hours at 4° C. Fixed placentas were weighed (wet weight, g) and converted into pre-embedding placental volumes (*V*_pre_) by dividing wet weight by tissue density (1.05 g/cm^3^) prior to multistage systematic and uniformly-random sampling (SURS) (Veras et al., 2008). Briefly, placentas were cut into ∼six 2-mm slices perpendicular to the chorionic plate. Slices were randomised to generate isotropic uniformly random (IUR) orientation and embedded in paraffin for stereological estimation of placental compartment volumes, maternal blood volume and surface area, and fetal capillary surface area. Shrinkage correction (performed per placenta). Shrinkage introduced during dehydration/clearing and paraffin embedding was quantified for each placenta individually by estimating the post-embedding placental volume (*V*_post_) using the Cavalieri Principle (Howard et al., 2005, Ribeiro et al., 2008). Each paraffin-embedded 2-mm slice was serially sectioned at 5 µm, and ∼10 evenly spaced sections per slice were selected systematic and uniformly-random through the block, mounted, and stained with Haematoxylin and Eosin (H&E). A point grid was applied to the sampled sections to estimate *V*_post_ by Cavalieri. The volume shrinkage coefficient for each placenta was calculated as:

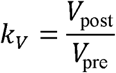

(with fractional shrinkage =1 - *k_V_*). Absolute compartment volumes estimated from paraffin sections were corrected back to pre-embedding dimensions as:

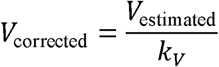

Surface area was estimated as the product of surface density and the corresponding shrinkage-corrected placenta volume:

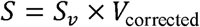

(Where S_V_ is the surface density estimated by intersection counting and v_corrected_ is the placenta volume corrected after the paraffin-embedding shrinkage.

### Placental RNA extraction and sequencing

Placental RNA was extracted from one quatre of an intact frozen placenta using the miRNeasy mini kit (Qiagen, UK), following tissue disruption in Qiazol (Qiagen, UK) using the Qiagen TissueLyserII (25Hz, 4x30 seconds), according to manufacturer’s instructions. RNA concentrations were assessed using Nanodrop and integrity was examined using Agilent 4200 TapeStation and the RNA ScreenTape Assay (Agilent; 5067-5576 and 5067-5577) with an acceptable RIN>8. Stranded RNA-seq libraries were prepared from 1000 ng of total RNA per sample, using the NEBNext Ultra II Directional RNA Library Preparation Kit for Illumina (NEB; E7760) and NEBNext Multiplex Oligos for Illumina (96 Unique Dual Index Pairs) (NEBNext; E6440). Prior to library preparation, total RNA was treated with QIAseq FastSelect -rRNA HMR, to prevent any rRNA present from being converted into sequencing library. Libraries were quantified using the Qubit Fluorometer and the Qubit dsDNA HS Kit (ThermoFisher Scientific; Q32854). Library fragment-length distributions were analysed using the Agilent 4200 TapeStation and the Agilent High Sensitivity D1000 ScreenTape Assay (Agilent; 5067-5584 and 5067-5585). Libraries were pooled in equimolar amounts and final library quantification was performed using the KAPA Library Quantification Kit for Illumina (Roche; KK4824). The library pool was sequenced on the Illlumina NextSeq500 over three NextSeq500 High Output 150 cycle kits (Illumina; 20024907), to generate over 40 million pairs of 75-bp paired-end reads per sample. Raw reads were trimmed of Illumina adapters and low quality (Q<20) nucleotides using TrimGalore (v 0.6.7) (https://www.bioinformatics.babraham.ac.uk/projects/trim_galore/). Reads shorter than 15 bp were discarded. Trimmed reads were aligned to Mus musculus reference genome GRCm39 using HISAT2 (v 2.2.1) (Kim et al., 2019). StringTie (v 2.2.1) (Kovaka et al., 2019) was used to assemble genes and calculate gene abundance. Differential expression analysis was performed using DESeq2. Data was visualised using ShinyGo, ClustVis, Network Analyst and Venny2.1 free web-based software using a pAdj<0.05, FDR <0.05 and no fold change cut off. All gene expression data is publicly available via Gene Expression Omnibus (GSE301011).

### Fetal sex determination

DNA was extracted from fetal tail tissue using the DNeasy Blood and Tissue kit (Qiagen, UK) according to the manufacturer’s instruction. For PCR determination of fetal sex, 2 μl (100 ng) of template DNA was added to a mastermix comprising 10 μl mastermix (2 × GoTaq® Green Master Mix, Promega UK), 1 μl primer mix (25 μM forward and reverse primers) and 5 μl water per reaction. Amplification was performed using an Techgene thermocycler using 3 primer sets specific for regions on the Y (Sry and Zfy) and X (Dxnds3) chromosomes (primer sequences are provided in Supplementary Table S2).

### Statistical Analysis

All data were assessed for normality with GraphPad Prism (version 10) or SPSS (version 28). Early placental (EPC) data were analysed using a One-way ANOVA for normally distributed data or a Kruskal-Wallis test for non-normally distributed data with appropriate post-hoc test. For the analysis of stud male growth, an additional FDR correction was applied to account for the multiple testing of weight at each individual week. Late gestation (E17.5) fetal and placental data were analysed using a generalised linear mixed model analysis with paternal origin of litter, and paternal duration on their respective diet incorporated as random effects (Watkins et al., 2018). Significance was taken at P <0.05.

## Author Contributions

Conceptualization: HLM, NE, NH, MC, SH, FS, MCU, IK, NN, RTM, FL, AAC, VB, AJW

Methodology: HLM, NE, NH, MC, SH, FS, MCU, IK, NN, RTM, FL, AAC, VB, AJW

Investigation: HLM, NE, NH, MC, SH, FS, MCU, IK, NN, AAC, VB, AJW

Data curation: HLM, NE, NH, MC, SH, FS, MCU, IK, NN, RTM, FL, AAC, VB, AJW

Formal analysis: HLM, NE, NH, MC, SH, FS, MCU, IK, NN, RTM, FL, AAC, VB, AJW

Funding acquisition: AJW, RTM

Writing – original draft: HLM, VB, AJW

Writing – review & editing, HLM, NE, NH, MC, SH, FS, VW, MCU, IK, NN, STM, RTM, FL, RSR, AAC, VB, AJW.

All authors have read and agreed to the published version of the manuscript.

## Data Availability Statement

All data are either contained within the Supplemental Materials or are available upon reasonable request sent to the corresponding author. Placental and testicular transcriptomic data have been deposited with in the Gene Expression Omnibus under accession numbers GSE301011 and GSE279868 respectively.

## Acknowledgments

The authors would like to thank the staff within the Bio Support Unit at the University of Nottingham for animal provision and maintenance.

## Figure legends

**Supplemental Figure 1:** Representative (A) no primary and (B) no secondary negative controls of testicular sections sained for Plzf and Ddx4. Representative (C) no primary and (D) no secondary negative controls of testicular sections sained for Sox9.

**Supplementary Table S1:**
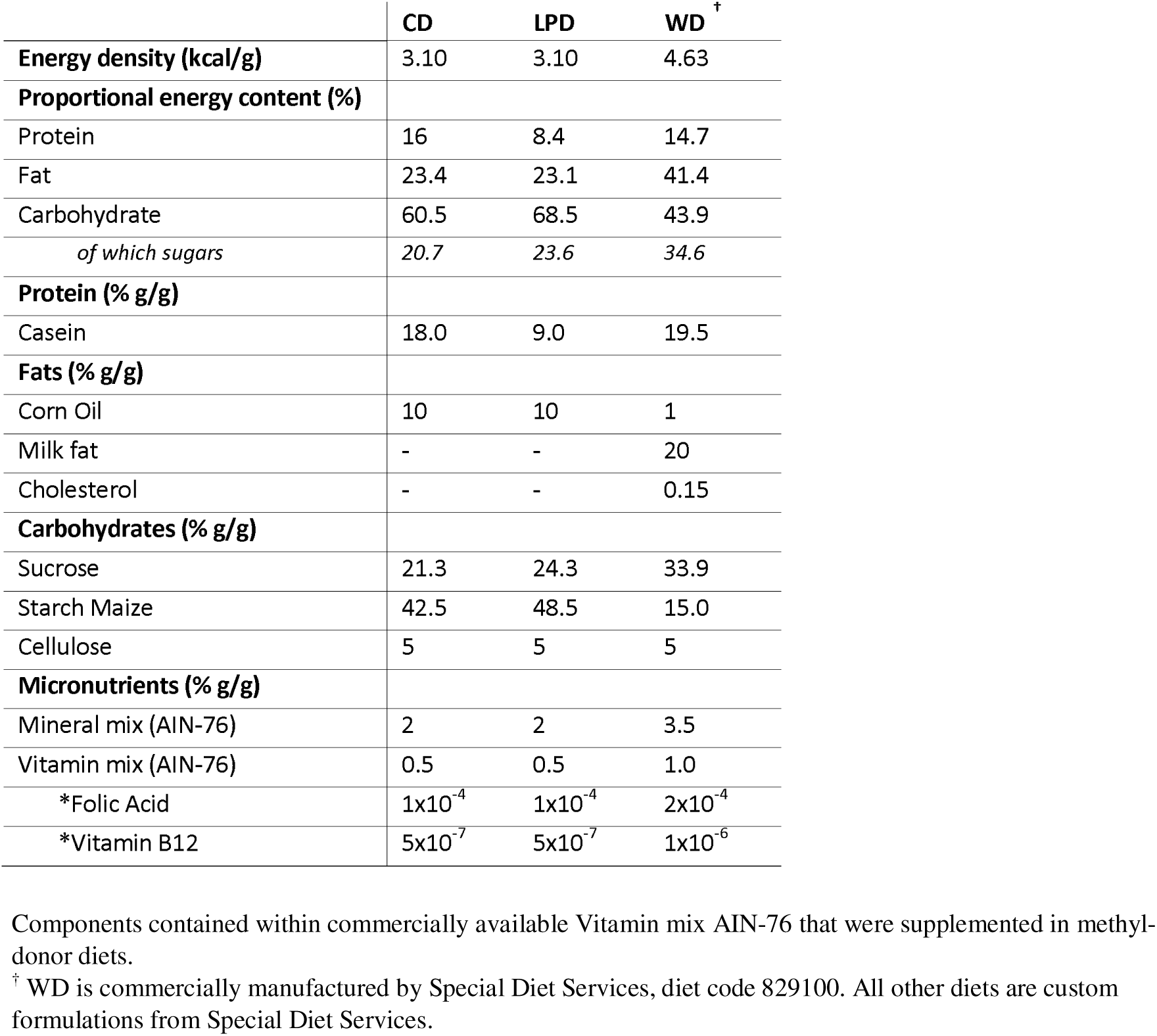
Ingredients and nutritional information of diets fed to male mice.

**Supplementary Table 2.**
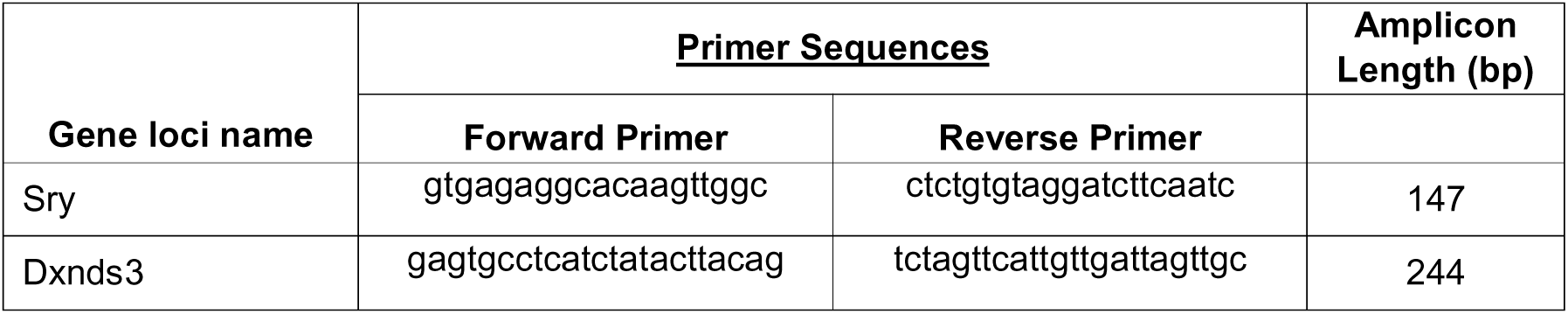
Sexing PCR primers.

