## Supplemental Table 1 for "Paternal over- and under-nutrition program fetal and placental development in a sex-specific manner in mice"

**Supplementary data**

**Table S1: Ingredients and nutritional information of diets fed to male mice**

|  | **CD** | **LPD** | **MD-LPD** | **WD^†^** | **MD-WD** |
| --- | --- | --- | --- | --- | --- |
| **Energy density (kcal/g)** | 3.10 | 3.10 | 3.10 | 4.63 | 4.63 |
| **Proportional energy content (%)** |  |  |  |  |  |
| Protein | 16 | 8.4 | 8.4 | 14.7 | 14.7 |
| Fat | 23.4 | 23.1 | 23.1 | 41.4 | 41.4 |
| Carbohydrate | 60.5 | 68.5 | 68.5 | 43.9 | 43.9 |
| *of which sugars* | *20.7* | *23.6* | *23.6* | *34.6* | *34.6* |
| **Protein (% g/g)** |  |  |  |  |  |
| Casein | 18.0 | 9.0 | 9.0 | 19.5 | 19.5 |
| **Fats (% g/g)** |  |  |  |  |  |
| Corn Oil | 10 | 10 | 10 | 1 | 1 |
| Milk fat | - | - | - | 20 | 20 |
| Cholesterol | - | - | - | 0.15 | 0.15 |
| **Carbohydrates (% g/g)** |  |  |  |  |  |
| Sucrose | 21.3 | 24.3 | 24.3 | 33.9 | 33.9 |
| Starch Maize | 42.5 | 48.5 | 46.0 | 15.0 | 12.3 |
| Cellulose | 5 | 5 | 5 | 5 | 5 |
| **Micronutrients (% g/g)** |  |  |  |  |  |
| Choline chloride | 0.2 | 0.2 | 0.7 | 0.2 | 0.7 |
| D,L-Methionine | 0.50 | 0.50 | 1.25 | 0.00 | 0.75 |
| Betaine | - | - | 1.5 | - | 1.5 |
| Mineral mix (AIN-76) | 2 | 2 | 2 | 3.5 | 3.5 |
| Vitamin mix (AIN-76) | 0.5 | 0.5 | 0.5 | 1.0 | 1.0 |
| *Folic Acid | 1x10^-4^ | 1x10^-4^ | 1.6x10^-3^ | 2x10^-4^ | 1.6x10^-6^ |
| *Vitamin B12 | 5x10^-7^ | 5x10^-7^ | 1.5x10^-4^ | 1x10^-6^ | 1.5x10^-4^ |

Components contained within commercially available Vitamin mix AIN-76 that were supplemented in methyl-donor diets.

^†^ WD is commercially manufactured by Special Diet Services, diet code 829100. All other diets are custom formulations from Special Diet Services.
