## Supplemental Table 2 for "Paternal over- and under-nutrition program fetal and placental development in a sex-specific manner in mice"

**Supplemental Table 2.** Sexing PCR primers.

| **Gene loci name** | **Primer Sequences** | | **Amplicon Length (bp)** |
| --- | --- | --- | --- |
|  | **Forward Primer** | **Reverse Primer** |  |
| Sry | gtgagaggcacaagttggc | ctctgtgtaggatcttcaatc | 147 |
| Dxnds3 | gagtgcctcatctatacttacag | tctagttcattgttgattagttgc | 244 |
